# Brainstem speech encoding is dynamically shaped online by fluctuations in cortical *α* state

**DOI:** 10.1101/2022.04.11.487894

**Authors:** Jesyin Lai, Caitlin N. Price, Gavin M. Bidelman

## Abstract

Experimental evidence in animals demonstrates cortical neurons innervate subcortex bilaterally to tune brainstem auditory coding. Yet, the role of the descending (corticofugal) auditory system in modulating earlier sound processing in humans during speech perception remains unclear. Here, we measured EEG activity as listeners performed speech identification tasks in different noise backgrounds designed to tax perceptual and attentional processing. Brainstem speech coding might be tied to attention and arousal states (indexed by cortical *α* power) that actively modulate the interplay of brainstem-cortical signal processing. When speech-evoked brainstem frequency-following responses (FFRs) were categorized according to cortical *α* states, low *α* FFRs in noise were found to be weaker, correlated positively with behavioral response times, more “decodable” via classifiers, and associated indirectly with other signal-in-noise perceptual performance. Our data provide evidence for online corticofugal interplay in humans and establish that brainstem sensory representations are continuously yoked to the ebb and flow of cortical states to dynamically update perceptual processing.

## 1. Introduction

Central auditory processing of target sounds is influenced by multiple factors, such as background noise (Binkhamis, Léger, Bell, Prendergast, O’Driscoll and Kluk, 2019; Billings, Gordon, McMillan, Gallun, Molis and Konrad-Martin, 2020; Song, Skoe, Banai and Kraus, 2011), task engagement (Saderi, Schwartz, Heller, Pennington and David, 2021; Shaheen, Slee and David, 2021), attention (Saiz-Alía, Forte and Reichenbach, 2019; Price and Bidelman, 2021), and arousal state (Mai, Schoof and Howell, 2019; Saderi et al., 2021). These factors may interact to modulate sound processing differentially. For instance, in complex listening environments, attention aids the selection of behaviorally-relevant inputs over irrelevant background noise to prioritize target cues for robust speech-in-noise (SIN) understanding (Price and Moncrieff, 2021). Attention is defined as a top-down cognitive process that alerts and orients listeners to focus on environmental and external stimuli (Petersen and Posner, 2012). There is ample experimental evidence in animals that cortical neurons innervate subcortex (i.e., inferior colliculus) (Beyerl, 1978; Bajo and Moore, 2005) to provide such top-down fine-tuning of subcortical auditory neural coding (Atiani, Elhilali, David, Fritz and Shamma, 2009; Gao and Suga, 2000; Suga and Ma, 2003). Attention has also been shown to influence all stages of human auditory processing from the inner ear to cortex (Galbraith, Olfman and Huffman, 2003; Giard, Collet, Bouchet and Pernier, 1994; Lukas, 1980; Picton and Hillyard, 1974; Rinne, Balk, Koistinen, Autti, Alho and Sams, 2008).

Still, while attentional modulation of auditory cortical activity is well established (Alho, Rinne, Herron and Woods, 2014; Alho and Vorobyev, 2007; Picton and Hillyard, 1974), attention effects on auditory brainstem activity in humans remains highly controversial (Galbraith and Kane, 1993; Varghese, Bharadwaj and Shinn-Cunningham, 2015; Dunlop, Webster and Simons, 1965; Picton, Hillyard, Galambos and Schiff, 1971). For instance, brainstem responses recorded during speech perception tasks typically fail to vary with listening state (Varghese et al., 2015), despite concomitant changes in cortex. In recent work, we used frequency-following responses (FFRs)–a scalp-recorded potential reflecting the brain’s spectrotemporal encoding of sound–to directly investigate attentional influences on auditory brainstem processing during speech listening tasks. Using source-reconstructed FFRs, we demonstrated attention enhances brainstem neural coding to aid noise-degraded speech perception (Price and Bidelman, 2021). We also found attentional enhancements in neural signaling between sub- and neo-cortical levels with stronger functional connectivity in the top-down (corticofugal) direction during challenging listening conditions. While these data provided neuroimaging evidence for efferent influences on brainstem auditory processing in humans, they did not speak to the temporal dynamics of auditory efferent system function. The allocation of attention is itself dynamic, waxing and waning over time (Kucyi, Hove, Esterman, Matthew Hutchison and Valera, 2017; Helfrich, Fiebelkorn, Szczepanski, Lin, Parvizi, Knight and Kastner, 2018). Presumably, changes in brainstem sound processing from corticofugal feedback occur online, as cortex dynamically shifts functional states dependent on stimulus or task demands (e.g., listener engagement, arousal, attention). Demonstrating FFRs depend on concurrent cortical activation would establish new and direct evidence that the corticofugal system dynamically shapes subcortical auditory processing in humans in real-time.

Besides attention, top-down pathways also regulate arousal (Krone, Frase, Piosczyk, Selhausen, Zittel, Jahn, Kuhn, Feige, Mainberger, Klöppel, Riemann, Spiegelhalder, Baglioni, Sterr and Nissen, 2017). While attention is usually directed towards external stimuli and involves the allocation of processing resources to relevant stimuli (Coull, 1998), arousal is related more to the internal state of physiological reactivity of the subject (Eysenck, 1982; Cohen, 2014; Robbins and Everitt, 1995), such as sleep, wakefulness, or excitement. When attending to sounds and actively engaging in a task, a subject’s internal arousal state is not static but fluctuates. Hence, we posited speech sound representations, as indexed via FFRs, might simultaneously track with trial-by-trial variations in cortical arousal state. Such findings would demonstrate online functional effects of the corticofugal pathways in the human auditory system.

In this study, we measured source-reconstructed brainstem and cortical neuroelectric electroencephalogram (EEG) activity as human listeners performed SIN tasks aimed to tax perceptual and attentional processing. Brainstem sound processing was measured via FFRs. FFRs represent sustained, phase-locked neural activity of a population of neurons which faithfully track dynamic sound features in the auditory brainstem (Bidelman, 2018; Smith, Marsh and Brown, 1975; White-Schwoch, Anderson, Krizman, Nicol and Kraus, 2019; Marsh, Worden and Smith, 1970; Coffey, Nicol, White-Schwoch, Chandrasekaran, Krizman, Skoe, Zatorre and Kraus, 2019; Tichko and Skoe, 2017). Internal arousal state is abstract and inherently subjective rendering it difficult to quantify behaviorally. Some EEG studies investigating arousal revealed that *α* suppression is induced by emotionally arousing stimuli (Uusberg, Uibo, Kreegipuu and Allik, 2013; Aftanas, Varlamov, Pavlov, Makhnev and Reva, 2002). As a result, we measured cortical *α* activity from listeners’ running EEG as they executed the speech perception tasks and used it as a neural proxy of arousal (Pivik and Harman, 1995). We hypothesized speech coding along the central auditory system might vary with attention and internal arousal states that actively modulate the interplay of brainstem-cortical signal processing. Furthermore, as *α* power is predictive of behavioral performance (Klatt, Getzmann, Begau and Schneider, 2020; Haegens, Nácher, Luna, Romo and Jensen, 2011; Kelly, Gomez-Ramirez and Foxe, 2009; Gould, Rushworth and Nobre, 2011), we explored correlations between *α*-yoked FFRs and speech perception measures. We also investigated neural decoding of FFRs at different

*α* power as this may provide further evidence revealing the role of *α* in being predictive of behavioral performance. Our results provide direct evidence that brainstem speech coding (particularly in noise) is modulated online so that sensory representations are continuously yoked to the ebb and flow of cortical states to dynamically update perceptual processing.

## 2. Methods and Materials

### 2.1. Participants

We recruited twenty young adults (age: 18–35 years, M = 24, SD = 3.4 years; 11 female). All participants exhibited normal hearing thresholds (≤ 25 dB HL; 250–8000 Hz). Since language background and music experience influence FFRs and SIN performance (Mankel and Bidelman, 2018; Parbery-Clark, Skoe and Kraus, 2009; Zhao and Kuhl, 2018), we required all participants to have < 3 years of formal musical training (M = 0.8 years, SD = 1.2) and speak native English. Participants were predominantly right-handed (M=82.04%, SD = 21.04) with no history of neuropsychiatric disorders. All provided written informed consent in accordance with the Declaration of Helsinki and a protocol approved by the University of Memphis IRB.

### 2.2. Speech stimuli and task

The speech stimuli and task are described fully in Price and Bidelman (2021) and illustrated in Fig. 1A. Brainstem and cortical EEG were recorded simultaneously (Bidelman, Moreno and Alain, 2013; Bidelman, Price, Shen, Arnott and Alain, 2019) during SIN-listening tasks to investigate the attentional effects and cortical influences on brainstem responses to speech. Three synthesized vowel tokens (e.g., /a/, /i/, /u/) were presented during the recording of EEGs. These vowels were chosen because they are sustained periodic sounds optimal for evoking FFRs (Bidelman et al., 2019; Skoe and Kraus, 2010). Each vowel was 100 ms with a common voice fundamental frequency (F0 = 150 Hz). The first and second formants were 730, 270, 300 Hz (F1) and 1090, 2290, 870 Hz (F2) for /a/, /i/, and /u/, respectively. Notably, the F0 of these stimuli is above the phase-locking limit of cortical neurons and observable FFRs in cortex (Bidelman, 2018; Brugge, Nourski, Oya, Reale, Kawasaki, Steinschneider and Howard, 2009), ensuring our FFRs would be of brainstem origin (Bidelman, 2018; Coffey, Herholz, Chepesiuk, Baillet and Zatorre, 2016). The vowels were presented in clean (i.e., no background noise) and noise-degraded conditions. For the noise condition, vowel stimuli were mixed with 8 talker noise babble (Killion, Niquette, Gudmundsen, Revit and Banerjee, 2004) at a signal-to-noise ratio (SNR) of +5 dB (speech at 75 dB_*A*_ SPL and noise at 70 dB_*A*_ SPL).

**Figure 1:**
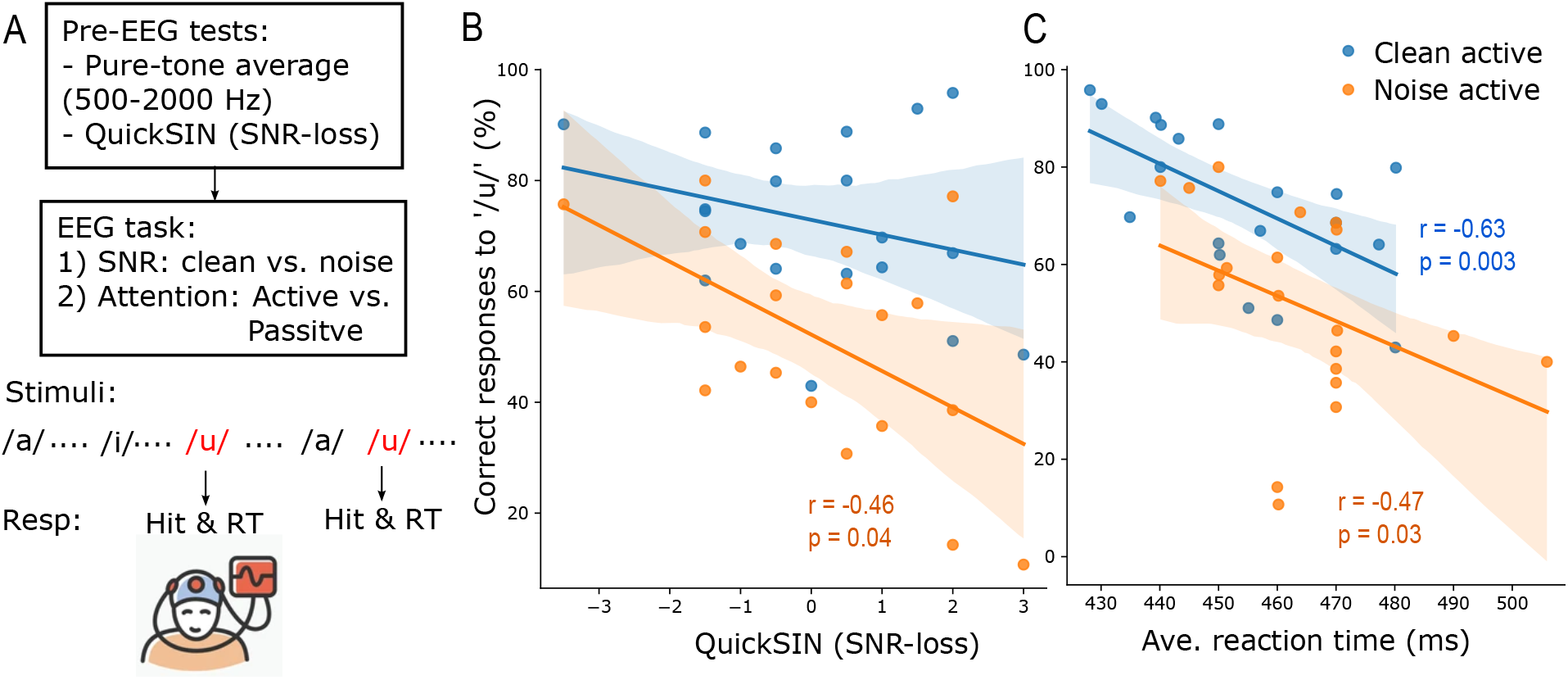
EEG task performance correlates with normative measures of SIN perception (i.e., QuickSIN scores). (A) Prior to EEG recordings, all participants’ pure-tone averages at 500-2000 Hz were obtained and speech-in-noise perception was assessed with QuickSIN. Subsequently, speech-EEGs were recorded under clean or noisy (+5 dB SNR) backgrounds and in active (/u/ detection task) or passive (no task) conditions. (B) QuickSIN scores, in which speech sentences were tested in noise, predict performance during the EEG task for noise-degraded (but not clean) speech. (C) In both clean and noise active sessions, speech detection accuracy predicts reaction times obtained during EEG tasks. r = Spearman’s correlation; shaded area = 95% CI of the regression line.

During active blocks, participants detected /u/ tokens, which were infrequent, via button press. A “hit” was defined as detection within 5 token presentations of /u/. For passive blocks, participants watched a captioned movie and were instructed to ignore any sounds they heard. This passive “task” has been shown to maintain arousal without impeding auditory processing (Pettigrew, Murdoch, Ponton, Kei, Chenery and Alku, 2004).

Each attentional block and noise condition (e.g., clean active, noise active, clean passive, noise passive) was divided into 2 runs allowing for breaks as needed. A single run lasted approximately 7.75 minutes and contained 1000 trials of each frequent token (/a/, /i/) and 70 trials of the infrequent /u/ token resulting in a total of 2070 trials per run (random order; jittered interstimulus = 95-155 ms, 5 ms steps, uniform distribution; rarefaction polarity) (cf. (Shiga, Althen, Cornella, Zarnowiec, Yabe and Escera, 2015)). Participants completed two runs of each attentional block and noise condition (e.g., active clean, active noise, passive clean, passive noise). Block (active/passive) and condition (clean/noisy speech) presentation were counterbalanced across participants to minimize order and fatigue effects (breaks were provided between blocks).

Stimulus presentation was controlled by MATLAB (The Mathworks, Inc.; Natick, MA) routed to a TDT RP2 interface (Tucker-Davis Technologies; Alachua, FL) and delivered binaurally through electromagnetically shielded (Campbell, Kerlin, Bishop and Miller, 2012) insert earphones (ER-2; Etymotic Research; Elk Grove Village, IL). Phantom bench tests confirmed this shielding fully eliminated stimulus electromagnetic artifact from contaminating EEG recordings (Price and Bidelman, 2021).

### 2.3. QuickSIN test

We used the Quick Speech-in-Noise (QuickSIN) test to assess listeners’ speech perception in noise (Killion et al., 2004). Listeners heard lists of 6 sentences, each with 5 target keywords spoken by a female talker embedded in four-talker babble noise. Target sentences were presented at 70 dB SPL (binaurally) at SNRs decreasing in 5 dB steps from 25 dB (relatively easy) to 0 dB (relatively difficult). SNR-loss scores reflect the difference between a participant’s SNR-50 (i.e., SNR required for 50 % keyword recall) and the average SNR threshold for normal hearing adults (i.e., 2 dB) (Killion et al., 2004). Higher scores indicate poorer SIN performance. Participants’ scores in this study ranged from -3.5 to 3 dB of SNR loss (M = 0.1, SD = 1.6), consistent with normal hearing.

### 2.4. EEG acquisition and preprocessing

EEGs were recorded from 64-channels at 10-10 electrode locations across the scalp (Oostenveld and Praamstra, 2001). Electrodes on the outer canthi and superior/inferior orbit monitored ocular artifacts. Impedances were ≤ 5 kΩ. EEGs were digitized at a high sample rate (5 kHz; DC—2000 Hz online filters; SynAmps RT amplifiers; Compumedics Neuroscan; Charlotte, NC) to recover both fast (FFR) and slow (cortical) frequency components of the EEG (Bidelman et al., 2013; Musacchia, Strait and Kraus, 2008).

EEG data were processed using Python 3.9.7. For sensor (channel-level) analyses, the data were re-referenced to the mastoid electrodes (M1 and M2). A common average reference was used for subsequent source FFR analysis. Ocular artifacts (saccades and blinks) were corrected in the continuous EEGs using independent component analysis (ICA; method=‘fastica’) (Flexer, Bauer, Pripfl and Dorffner, 2005; Hyvarinen, 1999). In this process, ICs best matching the pattern in the electrooculogram (EOG) were automatically identified and excluded from the data. Responses were then filtered from 130 to 1500 Hz [finite impulse response (FIR) filters; hamming window with 0.02 dB passband ripple and 53 dB stopband attenuation] to further emphasize brainstem activity (Bidelman et al., 2013; Musacchia et al., 2008).

### 2.5. Source FFRs

We transformed the cleaned sensor data (64-ch.) into source space using a virtual source montage (Scherg, Ille, Bornfleth and Berg, 2002). The source montage was comprised of a single regional dipole (i.e., current flow in x, y, z planes) positioned in the brainstem midbrain (i.e., inferior colliculus) [for details see Bidelman (2018); Price and Bidelman (2021); Bidelman and Momtaz (2021)]. Source current waveforms (SWF) from the brainstem dipole were achieved via matrix multiplication of the sensor data (FFR waveform) with the brainstem dipole leadfield (L) matrix (i.e., SWF = L^-1^ x FFR). This applied an optimized spatial filter to all electrodes that calculated their weighted contribution to the scalp-recorded FFRs in order to estimate source activity within the midbrain (Scherg and Ebersole, 1994; Scherg et al., 2002). This model explained >90 % of the scalp-recorded FFR (Price and Bidelman, 2021). We used only the z-oriented source activity given the predominantly vertical orientation of current flow in the auditory midbrain pathways relative to the scalp [x- and y-orientations contribute little to the FFR; Bidelman (2018)].

### 2.6. Cortical activities

Cortical *α* band activity was also extracted from the EEG and used as a running index of arousal state (high or low) during the speech listening task. *α* was quantified from the POz channel given the posterior scalp topography of *α* activity. To isolate *α*, we filtered recordings at 8-12 Hz (FIR filters). Filtered *α* activities were epoched with a time window of 195 ms (−50 to 145 ms in which 0 ms corresponded to the onset of an /a/ or /i/ token) to capture approximately 1-2 cycles of *α* band. Infrequent /u/ tokens were excluded from analysis due to their limited trials. We then measured the root mean square (RMS) amplitude of single trial *α* activity to quantify cortical arousal level over the duration of the perceptual task. RMS values were then normalized to the median of all RMS values of each run. Next, we visualized the distribution of trial-wise normalized *α* RMS via a histogram. Trials falling within the 0-25th percentile were categorized as low *α* power while those falling within the 75-100th percentile were categorized as high *α* power. This stratification resulted in ∼1000 trials each of low- or high-*α* power per task condition. We similarly measured cortical activity in different frequency bands (e.g., *β* band; 18-22 Hz) and electrode sites (Fz channel) as negative control analyses. *β* band is usually dominant at frontocentral scalp areas and is distal from our primary site for *α* quantification at the posterior (POz). Analyzing POz *β* and Fz *α* helped to rule out any unspecific/general effects of high or low EEG activity on FFRs.

### 2.7. Brainstem FFRs

We then derived FFRs according to the trial-by-trial cortical state, categorizing source FFRs based on whether *α* amplitude in the same epoch was either high vs. low. Source FFRs were then averaged for each *α* category, stimulus, condition, and subject. We then analyzed the steady-state portion (10–100 ms) of FFR waveforms using the FFT (Blackman window; 11.1 Hz frequency resolution) to capture the spectral composition of the response. F0 amplitude was quantified as the peak spectral maximum within an 11 Hz bin centered around 150 Hz (i.e., F0 of the stimuli). FFR F0 indexes voice pitch coding and has been shown to predict successful SIN perception (Mankel and Bidelman, 2018; Parbery-Clark et al., 2009). To compare F0 amplitudes of low- vs. high-*α*-indexed FFRs, we calculated an F0 ratio for each condition by dividing F0 amplitudes of high-*α*-indexed FFRs by those of low-*α*-indexed FFRs.

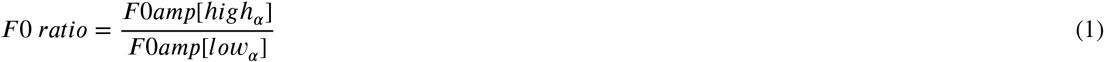

F0 ratios > 1 indicate brainstem FFRs were stronger during states of high cortical *α* power. In contrast, F0 ratios < 1 indicate FFRs were stronger during states of low cortical *α* power.

### 2.8. Statistical analysis

We used rmANOVAs to compare brainstem F0 ratios among the four conditions (clean active, noise active, clean passive, noise passive). Multiple pairwise comparisons (with Bonferonni corrections) were performed using the ‘pingouin’ package ins Python. One sample t-tests (‘scipy’ package in Python) were also used to evaluate whether FFR F0 ratios were significantly different from 1 (and thereby showed *α* modulation). To compare differences in *α* RMS values of all subjects across (clean vs. noise or active vs. passive) conditions, we performed post-hoc Conover’s test (‘scikit_posthocs’ package in Python), which is a non-parametric pairwise test, with Bonferroni adjustment. To test for differences in raw F0 amplitudes (log-transformed) across factors of our design, we used a 2 × 2 × 2 (POz *α*-band power x attention x SNR) mixed model (subjects = random factor) ANOVA (‘lme4’ package in Rstudio). This allowed us to compare raw F0 amplitudes of low- vs. high-*α*-indexed FFRs, clean vs. noise, and passive vs. active. Initial diagnostics were performed using residual and Q-Q plots to assess heteroscedasticity and normality of data. F0 amplitudes were log-transformed to improve normality and homogeneity of variance assumptions. Effect sizes are reported as η_*p*_^2^. We used Spearman’s correlations (‘scipy’ package in Python) to assess pairwise linear relations between neural and behavioral measures.

### 2.9. Decoding speech tokens from FFRs via machine learning

We used a linear SVM algorithm (Cristianini and Shawe-Taylor, 2000) to determine if the stimulus speech identity (i.e., /a/ vs. /i/ token) could be classified via low- and high-*α*-indexed FFRs. We focused on the noise active condition for this decoding analysis following procedures described by Xie, Reetzke and Chandrasekaran (2019). The epoched FFRs to /a/ or /i/ tokens were averaged, respectively, across ∼500 trials for each of the low- or high-*α*-indexed FFRs. These average FFR waveforms (10 to 100 ms steady state portion) were used as input features for SVM. Note this contrasts the F0 analyses in that the entire FFR spectrum was used for token-wise decoding. A total of 450 amplitude-by-time points were input as predictors; vowel types (i.e., /a/ & /i/) served as the ground truth class labels. During one iteration of training and testing, a four-fold cross-validation approach was used to train and evaluate the performance of the linear SVM classifier to obtain a mean decoding accuracy [Fig. 1 of Xie et al. (2019)]. In this process, subjects were randomly and equally divided into 4 subgroups with 5 unique subjects in each subgroup. Three of the 4 subgroups were selected as the training data while the remaining subgroup was used as the hold-out testing data. This was repeated within each iteration so that each subgroup was held-out as the test data whereas the other 3 subgroups were used to train the SVM classifier. Mean decoding accuracy (for distinguishing vowel tokens from FFRs) was calculated across cross-validated iterations. We performed a total of N=5000 iterations for each low and high *α* power.

To evaluate if the classifier accuracy (mean of N=5000 iterations) was statistically significant, ground truths (i.e., vowel types) were randomly assigned to FFR inputs, and the same training and testing procedures described above were repeated to derive a null distribution of decoding accuracies. We then calculated the p-value to determine the statistical significance of “true” classifier performance using the formula described in Phipson and Smyth (2010):

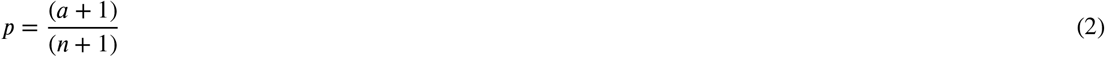

Where a is the number of decoding accuracies from the null distribution that exceeds the median of the actual distribution of decoding accuracies and n is the total number of decoding accuracies from the null distribution. The same p-value calculated using the above equation was also used to compare the prediction performance of the two “true” classifiers (low- vs. high-*α*-FFRs).

## 3. Results

### 3.1. Behavioral speech-in-noise performance

Speech-in-noise perception during the neuroimaging task was strongly associated with QuickSIN scores tested prior to EEG for noise-degraded but not clean speech (Spearman’s r = -0.46, p = 0.04, Fig. 1B). This confirms the external validity of our laboratory-based task in assessing clinically-normed SIN perception. Both perceptual outcomes were conducted in noisy backgrounds and under active listening conditions, suggesting token-wise vowel detection is a good proxy for sentence-level speech-in-noise perception. Behavioral hit responses (percent correct /u/ detections) were also negatively associated with response speeds in the EEG task regardless of SNR (clean active: r = -0.63, p = 0.003; noise active: r= -0.47, p = 0.03); slower decisions were associated with poorer accuracy (Fig. 1C).

### 3.2. Categorizing brainstem FFRs according to online cortical arousal state indexed by EEG *α* power

To determine if brainstem speech-FFRs are yoked to internal cortical arousal state, we measured trial-by-trial changes in EEG *α* band activity (∼10 Hz) to track low and high arousal states. At the same time, trialwise FFR F0 amplitudes were measured to index the magnitude of speech (i.e., voice-pitch) encoding at the brainstem level (Assmann, 1998; Mankel and Bidelman, 2018). FFRs were analyzed at the source level (see Methods) to unmix putative cortical contributions to the FFR (Coffey et al., 2016; Bidelman, 2018). We isolated cortical *α* band from the posterior POz channel, where waking *α* activity is maximal at the scalp (Moini and Piran, 2020; Nunez, 2016; Klatt et al., 2020). Unlike *α* band, *β* band is more prominent over frontcentral compared to posterior regions of cortex (Kropotov, 2009). To rule out any unspecific or general effects of cortical activity on FFRs, in addition to *α* band extracted from the POz channel (termed POz *α*), we also isolated *β* band from POz (termed POz *β*) and *α* from Fz (termed Fz *α*) to be used as negative controls.

Continuous POz *α* responses were epoched with a time window of -50 to 145 ms with respect to vowel onset time (i.e., the same analysis window as FFRs). We then computed the root-mean-square (RMS) power within each epoch. This resulted in approximately 4000 trials of POz *α* RMS values (Fig. 2A) corresponding to the 4000 frequent speech tokens in the task (i.e., /a/ and /i/). All RMS values were normalized to the RMS median of each run and visualized using a histogram (Fig. 2B). Subsequently, RMS values falling within the range of 0-25th percentile were categorized as low *α* power while RMS values falling within 75-100th percentile were considered as high *α* power. This resulted in ∼1000 trials in each of the lower and upper percentile ranges. Trials outside these ranges were not analyzed because we wanted to maximize differences between low- and high-*α*-indexed FFRs.

**Figure 2:**
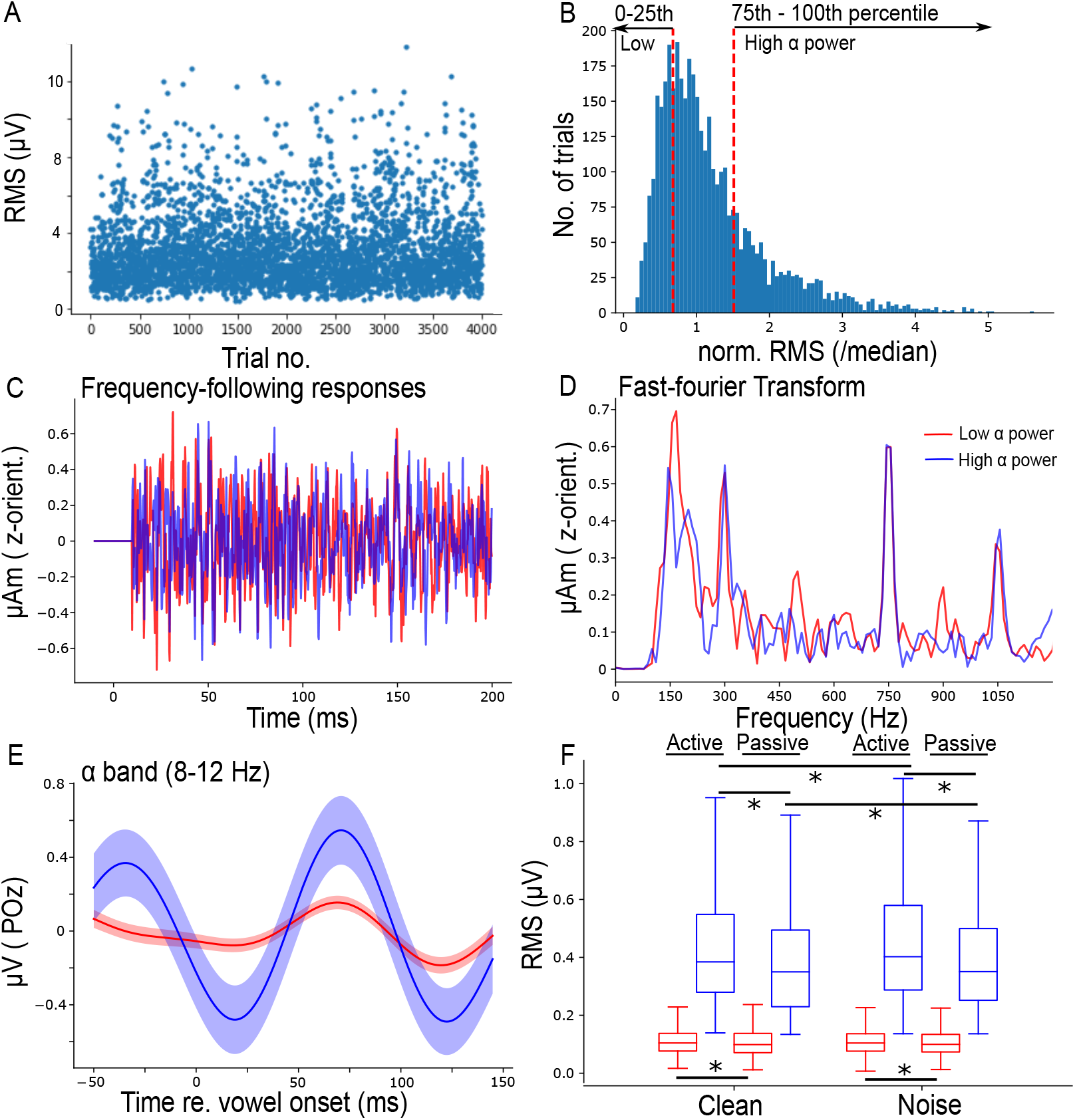
Examples of data processing (representative subject) to separate FFRs based on cortical arousal state (i.e., *α* power). (A) Cortical *α* band was isolated from the POz channel and epoched at -50-145 ms with respect to vowel stimulus onset time. Root-mean-square (RMS) value was computed for every epoch (i.e., approximately 4000 single trials) per listener. (B) Histogram of RMS values normalized to the RMS median of each run. The first red dashed line (left) indicates the 25th percentile and the second (right) indicates the 75th percentile. RMS values falling within the range of 0-25th and 75-100th percentile were categorized as low vs. high *α* power, respectively, yielding 1000 trials per state. (C) Average source FFRs (with removed onset responses) during low (red) vs. high (blue) EEG *α* power. (D) Frequency spectra of the steady state (10-100 ms) portion of FFR waveforms. (E) Average *α* waveform of low and high power for the trials extracted from the lower or upper 25th percentiles in (B). (F) Boxplots showing RMS values of low and high *α* power of all subjects (N=20). Shaded area = ± s.e.m. * p< 0.01 (Conover’s test, non-parametric pairwise test, with Bonferroni adjustment)

Speech-evoked source FFRs were then categorized based on their corresponding power in the same epoch window esulting in a set of low- vs. high-*α*-indexed FFRs. FFR time waveforms and spectra for a representative subject are shown in Fig. 2C and Fig. 2D for low- and high-*α* states. Corresponding average *α* responses are shown in Fig. 2E. The difference in activity level of low vs. high *α* power is apparent. RMS values of low and high *α* across all subjects for each of the four conditions is shown in Fig. 2F, with statistically significant differences (p <0.01) observed between some conditions of clean vs. noise or active vs. passive RMS values (non-parametric post-hoc Conover’s test with Bonferroni adjustment). The same process of categorizing the level of cortical bands and FFRs into low or high level was repeated on POz *β* and Fz *α* as well. These negative controls allowed us to compare the subsequent observations from POZ *α* to POZ *β* and Fz *α*, to ensure the observed changes in speech-evoked FFRs were specifically associated with cortical arousal level (indexed by *α* power) rather than general fluctuations in the EEG, per se.

### 3.3. Speech FFRs depend on SNR, attention, and cortical *α* power

Grand averaged FFR spectra for low- and high-*α*-indexed FFRs across conditions are shown in Fig. 3A-D. In noise, FFR F0 amplitudes during low *α* power were lower than F0 amplitudes during high *α* power, especially in the passive condition where low vs. high *α* F0 responses were significantly different (t = 2.18, p = 0.04). We computed a normalized (within-subject) measure of *α* FFR enhancement by dividing F0 amplitude of high-*α*-indexed FFRs by F0 amplitudes of low-*α*-indexed FFRs (“F0 ratio”). F0 ratios in noise were higher than in clean backgrounds (rmANOVA, F = 4.84, p = 0.04,η_*p*_^2^ = 0.20) (Fig. 3E). However, responses were comparable in the negative control (POZ *β* and FZ *α*) analyses, suggesting FFR modulations were specific to the cortical *α* band (indexing arousal) and not general properties of the EEG, per se (Fig. A.1). A mixed-model ANOVA on log F0 amplitudes also revealed significant main effects of SNR (clean vs. noise), attention (active vs. passive), and *α* power (low vs. high) (see Fig. 3F and Table 1).

**Figure 3:**
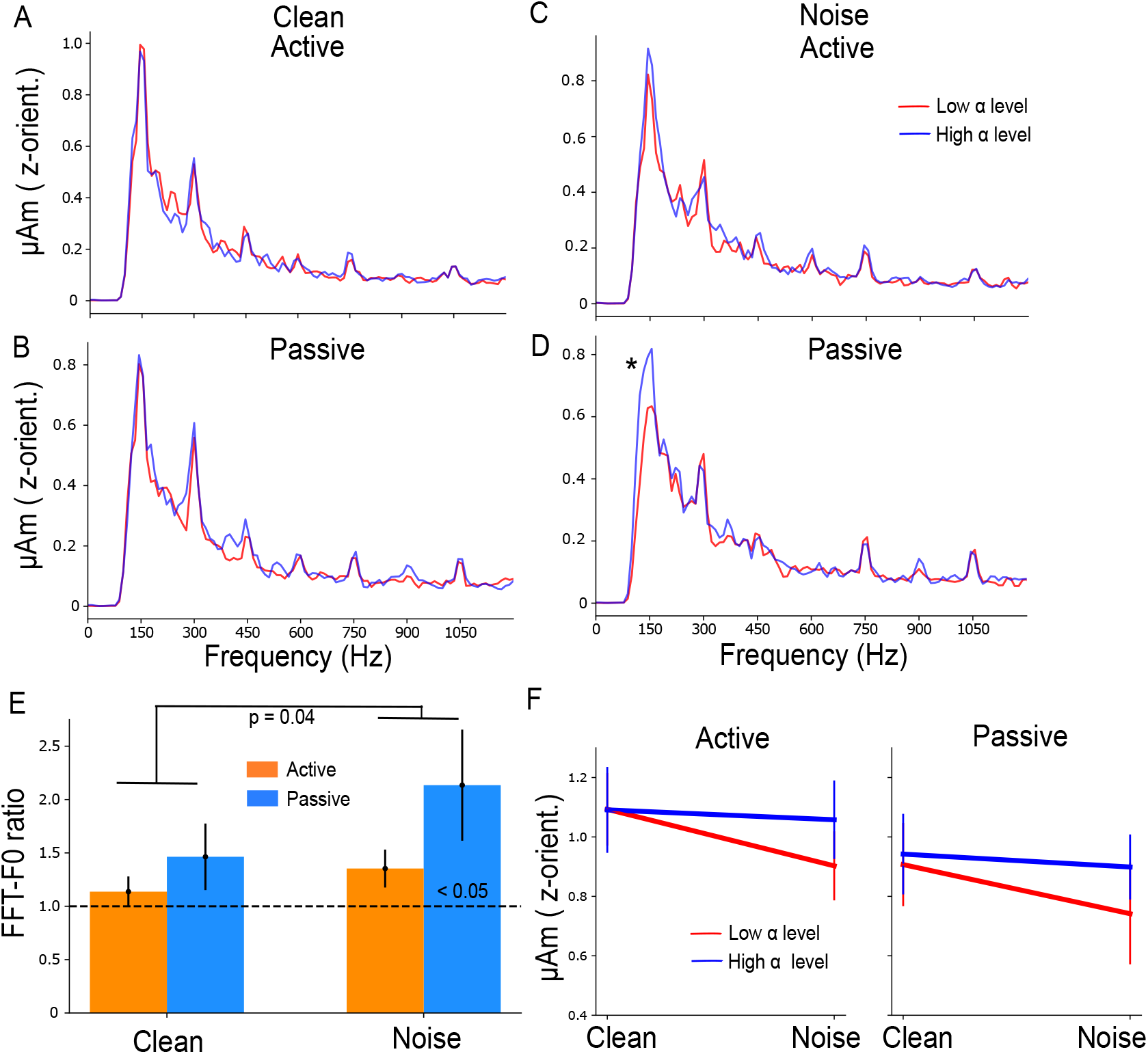
FFR F0 amplitudes during low *α* are smaller than during high *α* in noisy (but not in clean) backgrounds. (A-D) Frequency spectra of speech-evoked FFRs illustrating smaller F0 amplitudes during low *α* in both active and passive noisy backgrounds (C & D) but not in clean backgrounds (A & B). (E) A noise effect is observed for F0 ratio. Bars marked (<0.05) are significantly larger than 1 (1-sample t-test). (F) Raw FFR F0 amplitudes as a function of SNR (clean vs. noise), attention (active vs. passive) and *α* power (low vs. high). Errorbars= ± s.e.m., *p<0.05

**Table 1:** Results of Type III, mixed-model ANOVA on ln(FFR F0 amplitudes).

| | NumDF | DenDF | F-value | p-value | $\eta_p^2$ |
| --- | --- | --- | --- | --- | --- |
| SNR | 1 | 140 | 4.156 | 0.043 * | 0.03 |
| attention | 1 | 140 | 13.003 | <0.001 *** | 0.08 |
| $\alpha$ power | 1 | 140 | 5.027 | 0.027 * | 0.03 |
| SNR x attention | 1 | 140 | 0.136 | 0.73 | $9.71 \times 10^{-4}$ |
| SNR x $\alpha$ power | 1 | 140 | 2.248 | 0.136 | 0.02 |
| attention x $\alpha$ power | 1 | 140 | 2.008 | 0.158 | 0.01 |
| SNR x attention x $\alpha$ power | 1 | 140 | 0.166 | 0.685 | $1.18 \times 10^{-3}$ |

\* $p < 0.05$ , \*\*\* $p < 0.001$

### 3.4. Brain-behavior relations

We next assessed associations between behavioral performance in the speech detection task collected during EEG recordings (percent correct, RTs) and neural measures. The correlations of these behavioral metrics with FFR F0 amplitudes (per condition) were then tested systematically. Correlations for the clean condition were not significant. In contrast, for noise, we found a trend whereby in states of high cortical *α*, FFR strength in noise decreased for trials with slower RTs (combining both active and passive conditions, Fig. 4A). When focusing on the noise active condition, FFRs during low *α* states were strongly associated with RT (Spearman’s r = 0.54, p = 0.02). The brain-behavior relation between RTs and FFR amplitudes also differed as a function of cortical *α* power (Fisher’s z = 3.13, p = 0.001). This opposite direction in correlations was not observed between noise passive FFRs and RTs (recorded in the active task) (Fig. 4C). Repeating the same correlation analysis for POz *β* and Fz *α* in the noise active condition did not reveal any significant association between FFRs and RT (Fig. A.2). This suggests that the observed correlation between FFRs and RT was specifically related to *α* power in POz channel.

**Figure 4:**
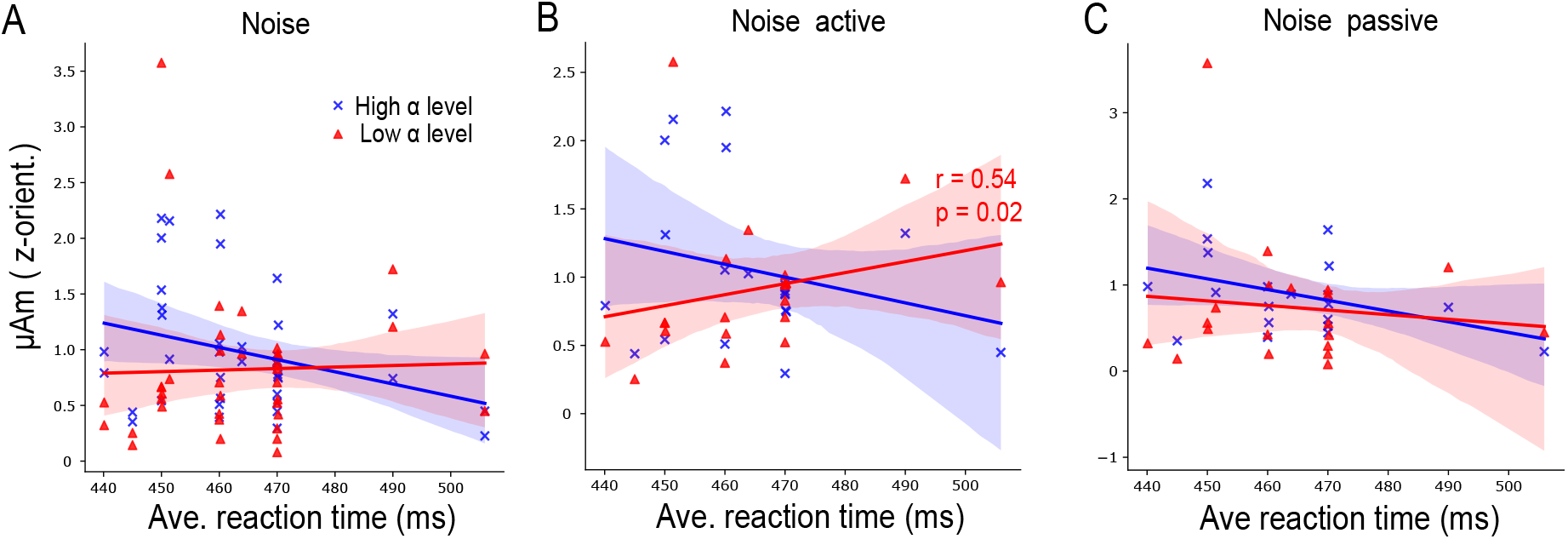
Brain-behavior associations between FFR and speech perception depend on cortical state. (A) In noise (pooling active and passive conditions), there is a trend of decreased FFR F0 amplitudes during high *α* power in listeners with slower reaction times. (B-C) When correlations were analyzed separately for noise active (B) and passive (C) conditions, FFRs during low *α* power correlated with reaction times in the active, but not the passive, condition. r = Spearman’s correlation, shaded area indicates 95 % confidence interval of the regression line.

### 3.5. Token decoding from FFRs during low & high *α* power

The previous analyses showed low-*α*-indexed FFR amplitudes were correlated with behavior in the noise active condition. We further asked whether FFRs during low *α* states are actually better at decoding speech representations (i.e., /a/ vs. /i/ tokens) compared to their high *α* counterparts. We followed the machine learning (ML)-based approach reported by Xie et al. (2019) to analyze FFRs for token decoding. Two separate linear support vector machine (SVM) classifiers were trained and tested on FFR waveforms with the goal of decoding the speech stimulus from neural responses. Mean linear SVM classification accuracies using low- or high-*α*-indexed FFRs are shown as confusion matrices in Fig. 5A. All FFRs yielded classification performance significantly above chance (Fig. 5B), confirming brainstem neural representations closely mirror the spectrotemporal properties of speech. There was more than a 20% increase (78.34 vs 57.77%) in the mean classification accuracy for /a/ tokens when using low-*α*-FFRs as inputs. The mean classification accuracies for /i/ tokens were at ceiling and exceeded 90% for FFRs at low (96.90%) and high (93.87%) *α* power. Furthermore, the classifier performance for low *α* was also significantly better than high *α* (Fig. 5B). These decoding findings suggest FFR speech representations were of higher fidelity (discriminability) under low compared to high *α* cortical states. Better vowel decoding may contribute to enhanced token discrimination in noisy backgrounds.

**Figure 5:**
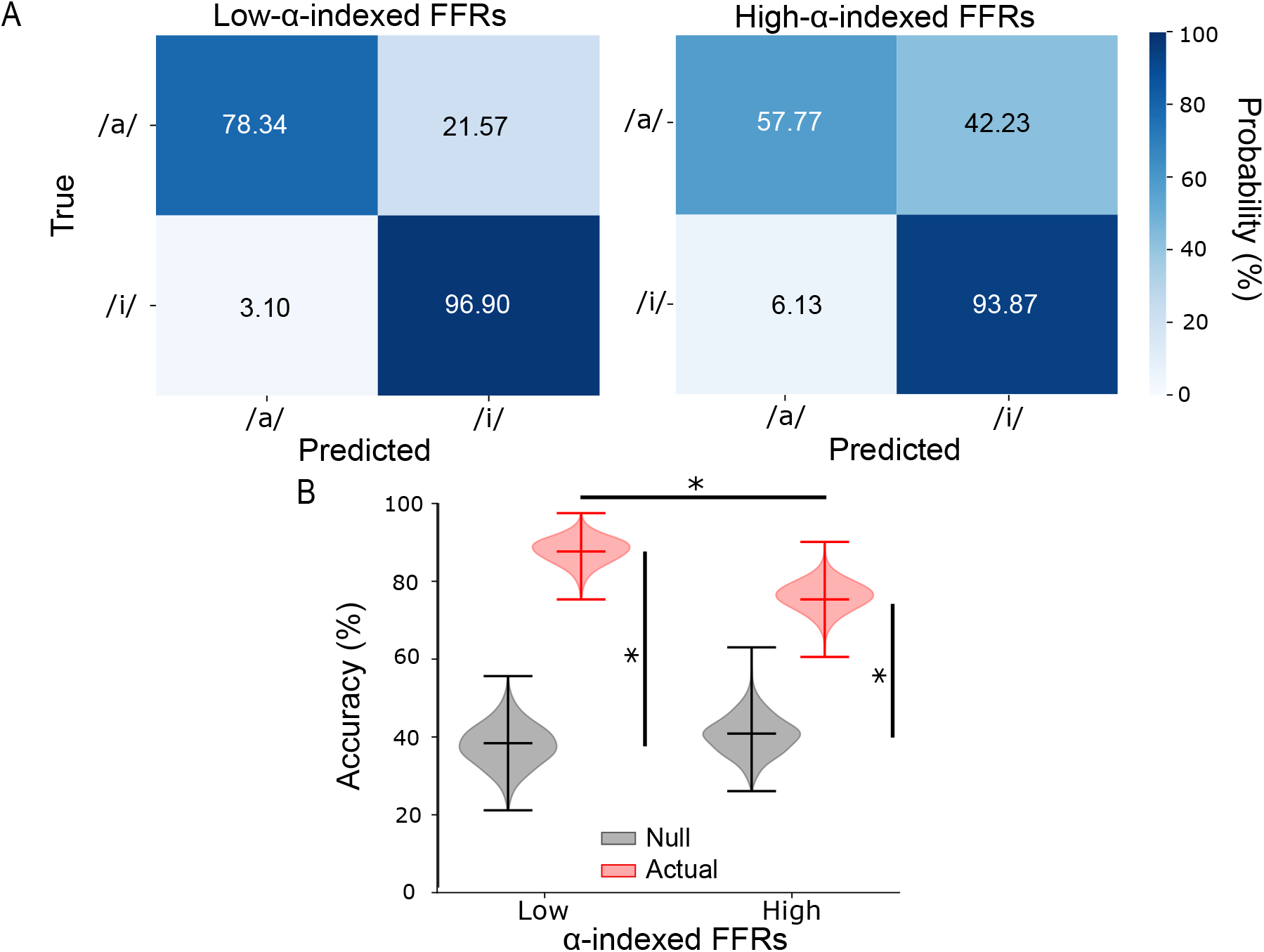
Average accuracy of linear SVMs in classifying /a/ and /i/ tokens (noise active condition) is better when using low-*α*-indexed FFRs as inputs. (A) Vowel decoding confusion matrices for low- (left) vs. high (right) *α*-indexed FFRs (N=5000 bootstrap iterations). (B) Overall distributions of prediction accuracies of SVM classifiers were significantly better using FFRs at low compared to high *α* power. Distribution of prediction accuracies for FFRs (actual) were also significantly above the null prediction accuracies. Upper/lower ticks=max/min; center tick=medians. * p< 0.001

## 4. Discussion

Our previous study revealed strong attentional enhancements in speech-FFRs at the source level and top-down neural communication from primary auditory cortex to brainstem that aids SIN perception (Price and Bidelman, 2021). Building upon those findings, we show here the existence of active and dynamic modulation of brainstem speech processing dependent on online changes in the listener’s cortical state. We found higher cortical *α* states positively associate with larger speech-evoked FFRs in adverse listening conditions. Furthermore, FFRs during low *α* power and recorded in the noise active condition predicted behavioral RTs for rapid speech detection and were associated indirectly with other perceptual measures of SIN performance. This positive relationship of FFRs and behavior at low *α* power were further advocated by better token classification using a neural decoding approach. As a whole, these data help reconcile equivocal findings by revealing top-down cortical activity dynamically influences brainstem encoding of speech, which directly imposes constraints on behavior.

Attention effects on brainstem responses have been highly controversial. Some studies reveal attention increases the robustness and temporal precision of speech-evoked FFRs (Galbraith et al., 2003; Forte, Etard and Reichenbach, 2017; Hartmann and Weisz, 2019; Lehmann and Schönwiesner, 2014). However, other studies have observed mixed (Holmes, Purcell, Carlyon, Gockel and Johnsrude, 2018; Saiz-Alía et al., 2019) or no attention effects on FFRs (Galbraith and Kane, 1993; Varghese et al., 2015). Attention effects at the brainstem level could be too subtle to detect in scalp EEG given the inherent mixing of intracranial sources (Vollmer, Beitel, Schreiner and Leake, 2017; Price and Bidelman, 2021) and/or require more difficult perceptual tasks (e.g., SIN perception) that challenge speech processing (Lehmann and Schönwiesner, 2014; Galbraith and Arroyo, 1993). In this study, we observed overall enhanced FFRs regardless of the background SNR when subjects actively engaged in challenging speech listening tasks (Fig. 3F, left panel). However, FFR changes due to cortical *α* were more prominent in noise than clean conditions (Fig. 3E-F). Similarly, even under passive listening conditions, these cortical dependencies in the FFR were stronger under states of low *α* and for noisy speech (Fig. 3E & right panel of F). These findings suggest that cortical influences like attention (Price and Bidelman, 2021) and internal arousal state (present study) on the FFR are more prominent in adverse listening conditions.

Elevated *α* oscillations were initially thought to reflect states of wakefulness but without active engagement in tasks (Pfurtscheller, Stancák and Neuper, 1996). Hence, in this study, we interpret low *α* power to indicate subjects were in a high arousal state and high *α* power to index task focus but in a state of wakeful relaxation. In both active and passive noise conditions, we observed larger speech-evoked FFRs during high *α* power (i.e., subjects in a relaxed, low arousal state). Our observations do not coincide with the findings of Mai et al. (2019) in which they reported higher phase-locked responses to speech at sub-cortical levels were associated with high arousal state. However, that study recorded EEGs during sleep and not during active tasks, and FFRs were separated according to sleep spindles to index subjects’ arousal states rather than online *α* as done here. These differences may contribute to the discrepancy between studies. Moreover, Makov, Sharon, Ding, Ben-Shachar, Nir and Golumbic (2017) reported greater phase-locked FFRs in wakefulness compared to sleep. Although stimuli are processed by the brain during sleep (Issa and Wang, 2008; Nir, Vyazovskiy, Cirelli, Banks and Tononi, 2015), neural responses to speech in the auditory subcortex reduce during sleep compared to wakefulness (Portas, Krakow, Allen, Josephs, Armony and Frith, 2000). Since EEG recordings were conducted in subjects during wakefulness in this study, our results may suggest that subcortical auditory processing in low vs. high arousal states are different in wake vs. sleep states.

In addition, *α* oscillations play a significant role in functionally inhibiting the processing of task-irrelevant information (Jensen and Mazaheri, 2010; Foxe and Snyder, 2011). Both *α* power and phase were found to modulate neuronal spike rate (Haegens et al., 2011) and thus can directly affect the efficiency of neural information flow. For example, Wilsch, Henry, Herrmann, Maess and Obleser (2015) demonstrated increased *α* power when participants listened to stimuli presented in noisy backgrounds. Induced *α* activity is crucial for speech processing in challenging listening conditions as it suppresses irrelevant information (Strauß, Wöstmann and Obleser, 2014). Here, we demonstrate that *α* power is significantly higher (p < 0.01, Conover’s test) in active than passive speech listening conditions and the mean RMS values of high *α* power is larger in the noise compared to the clean active condition (Fig. 2F). Another interesting and crucial point regarding *α* activity is that it fluctuates in amplitude with stimulus and task demands (Klimesch, 2012). Increased *α* may reflect inhibition to de-prioritize distractors while decreased *α* might reflect a release from inhibition to prioritize targets. Indeed, *α* oscillations have been interpreted to have two mechanistic roles underlying the functions of selective attention (Klimesch, 2012; Klatt et al., 2020). Klatt et al. (2020) were able to tease apart these two mechanisms by showing a shifting of attention towards lateralized targets resulted in contralateral decreases in *α* power whereas shifting attention away from lateralized distractors resulted in contralateral increases in *α* power. Furthermore, decreased *α* power has been associated with increased neural firing (Haegens et al., 2011) and improved behavioral performance (Haegens et al., 2011; Kelly et al., 2009; Gould et al., 2011).

In this vein, we observed listeners with smaller amplitudes of low-*α*-indexed FFRs having shorter behavioral RTs in the noise active condition (Fig. 4B). This highlights a relationship between low *α* power and behavioral performance. In the vowel identification task in noise, subjects were required to press a button whenever /u/ was presented. Hence, /u/ is a task-relevant target, background noise is task-irrelevant information while /a/ and /i/ can be considered as task-relevant distractors. Although we were unable to measure FFRs to /u/ due to insufficient token counts (only 140 trials in total), we speculate there would be increased neural firing to /u/ (task-relevant target) but lower neural responses to /a/ and /i/ (task-relevant distractors) during low *α* power. On the other hand, task-irrelevant background noise was inhibited during high *α* power but not /a/ and /i/ since they are task-relevant stimuli. These two mechanisms provide a possible explanation for our observations in the noise active condition where subjects with faster RTs had larger high-*α*-indexed FFR amplitudes and smaller low-*α*-indexed FFR amplitudes (Fig. 4B). Reversing these phenomena resulted in slower RTs when participants were actively identifying stimulus targets in noise.

At this stage, we have evidence showing FFR F0 amplitudes during low and high *α* power were associated with RT in the noise active condition. Moreover, RT was found to be strongly correlated to the percent of correct responses to /u/ (Fig. 1C) and percent of correct responses to /u/ obtained in noise was strongly predictive of QuickSIN scores (Fig. 1B). Collectively, these results show FFRs during low *α* power have an indirect relationship with other perceptual performance (e.g., percent of correct responses to /u/ and QuickSIN) tested in noise (Fig. A.3). In other words, cortical *α* modulation of FFRs in noisy backgrounds is indirectly associated with speech detection performance and QuickSIN via behavioral RT.

To further assess the behavioral relevance of *α*-FFR modulations to SIN listening, we investigated vowel decoding of FFRs at low vs. high *α* states. Similar ML approaches have been applied to speech-evoked FFRs to decode stimulus classes, e.g., Mandarin lexical tones (Llanos, Xie and Chandrasekaran, 2017; Xie et al., 2019) and speech tokens (Xie, Girshick, Dollár, Tu and He, 2017; Cheng, Xu, Gold and Smith, 2021). In these studies, decoding performance in correctly classifying FFRs is used as an objective measure of speech discrimination. Moreover, Xie et al. (2017) and Cheng et al. (2021) demonstrated that training and attention can improve FFR classification. The logic behind these ML-based approaches is that attention or training-related plasticity on auditory neural responses produce direct changes in the accuracy with which FFRs are classified. An increase in classification accuracy reflects positive modulation (i.e., enhanced or more robust neural representations to stimuli) while a decrease in accuracy implies negative modulation (i.e., reduced or less robust neural responses). The enhanced classification accuracies for /a/ and /i/ tokens we find under low-*α* EEG suggests speech representations are of higher fidelity under high arousal states. This higher fidelity could result from better response SNR or more consistent responses across subjects in the low-*α* state. This may also account for why brain-behavior relations between FFRs and RTs are mainly observed during low *α* states (Fig. 4B).

More broadly, difficulties in SIN perception (Duquesnoy, 1983; Helfer and Wilber, 1990; Fostick, Ben-Artzi and Babkoff, 2013) and changes in brainstem auditory processing in the presence of noise maskers have been reported in older listeners, even when stimulus intensities are matched for audibility (Lai and Bartlett, 2018). Meanwhile, some previous studies found age has an effect on brain oscillatory activity of *α* band (Yordanova, Kolev and Başar, 1998; Klimesch, 1999; Böttger, Herrmann and Von Cramon, 2002). Therefore, *α* oscillations in demanding and challenging listening tasks might be used as an indicator of age-dependent auditory cognitive effort of noise inhibition. Investigating *α* power and SIN perception in older adults may reveal how aging impacts top–down attentional control to facilitate processing of task-relevant target sounds and inhibit processing of task-irrelevant distractors or maskers. This could partly explain why older listeners have difficulties participating in cocktail party-like listening situations compared with younger listeners (Pichora-Fuller, 2003).

## 5. Conclusion

Our novel findings reveal that human brainstem FFRs to speech are dynamically modulated by cortical activity (indexed by *α* power) during speech-in-noise perception. FFRs are thus yoked online to shifts in internal arousal state. Our results also show FFRs during low *α* (i.e., high arousal state) have more “decodable” speech representations and are more predictive of behavioral SIN performance. Our novel paradigm and single-trial analysis of the FFR help resolve ongoing debates regarding attentional and/or arousal influences on brainstem auditory encoding and corticofugal engagement during active SIN perception in humans. Our data also elucidate possible mechanisms concerning the function of cortical *α* during SIN recognition whereby sound targets are actively segregated from task-relevant distractors and task-irrelevant background noise to improve perceptual outcomes.

## Data and code availability statement

The data supporting the reported findings are available from the corresponding author upon reasonable request.

## Ethics statement

All participants provided written informed consent prior to participation in accordance with protocols approved by the University of Memphis IRB.

## Acknowledgments

This work was supported by the National Institutes of Health (NIH/NIDCD R01DC016267) (G.M.B.).

## A. Appendix

**Figure A.1:**
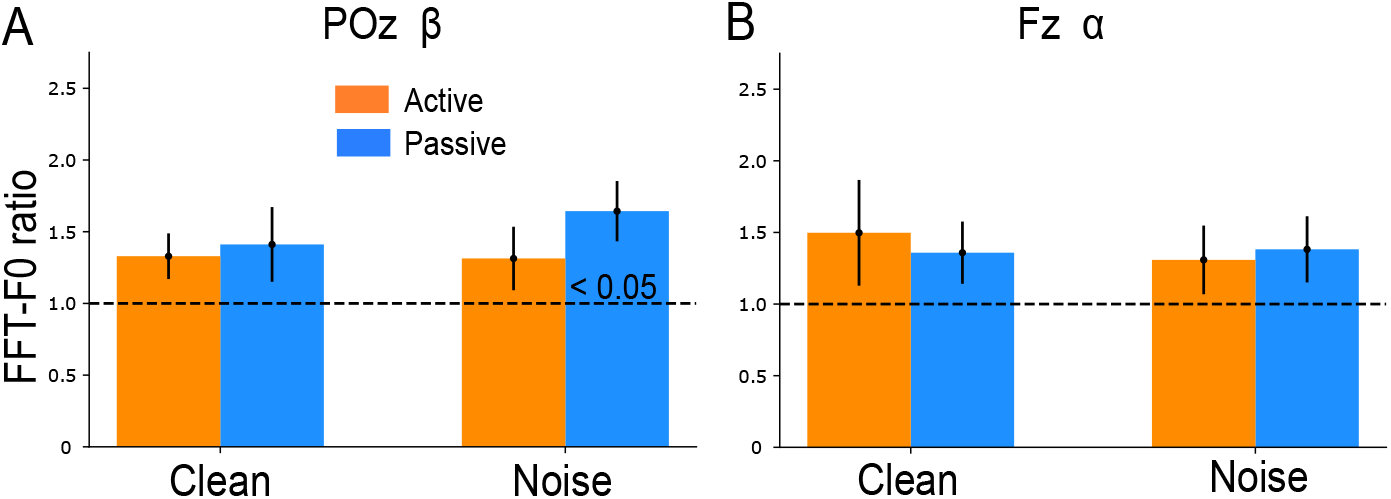
These control analyses showed no significant dependence of the FFR (as in Fig. 3E but for POz *β* band and Fz *α* band). No significant noise effect is observed in F0 ratio when comparing F0 amplitudes of high to low *β* in POz channel (A) or *α* in Fz channel (B). <0.05 in bars indicates significantly larger than 1 (1-sample t-test).

**Figure A.2:**
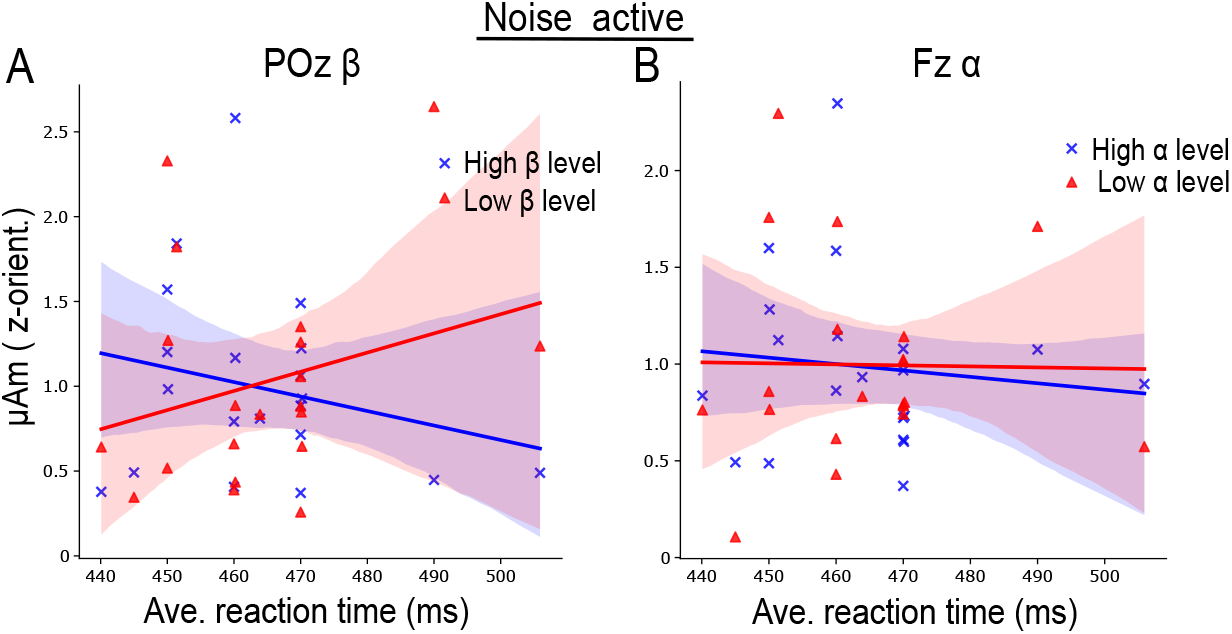
Repeating the same correlation analysis with a different EEG frequency band (*β*) and electrode (Fz) produced no significant result. No significant association is observed between F0 amplitudes of FFRs at low (red line) or high (blue line) cortical activity level and reaction time when analyzing *β* band from POz channel (A) or *α* band from Fz channel (B). Shaded area indicates 95 % confidence intervals of the regression line.

**Figure A.3:**
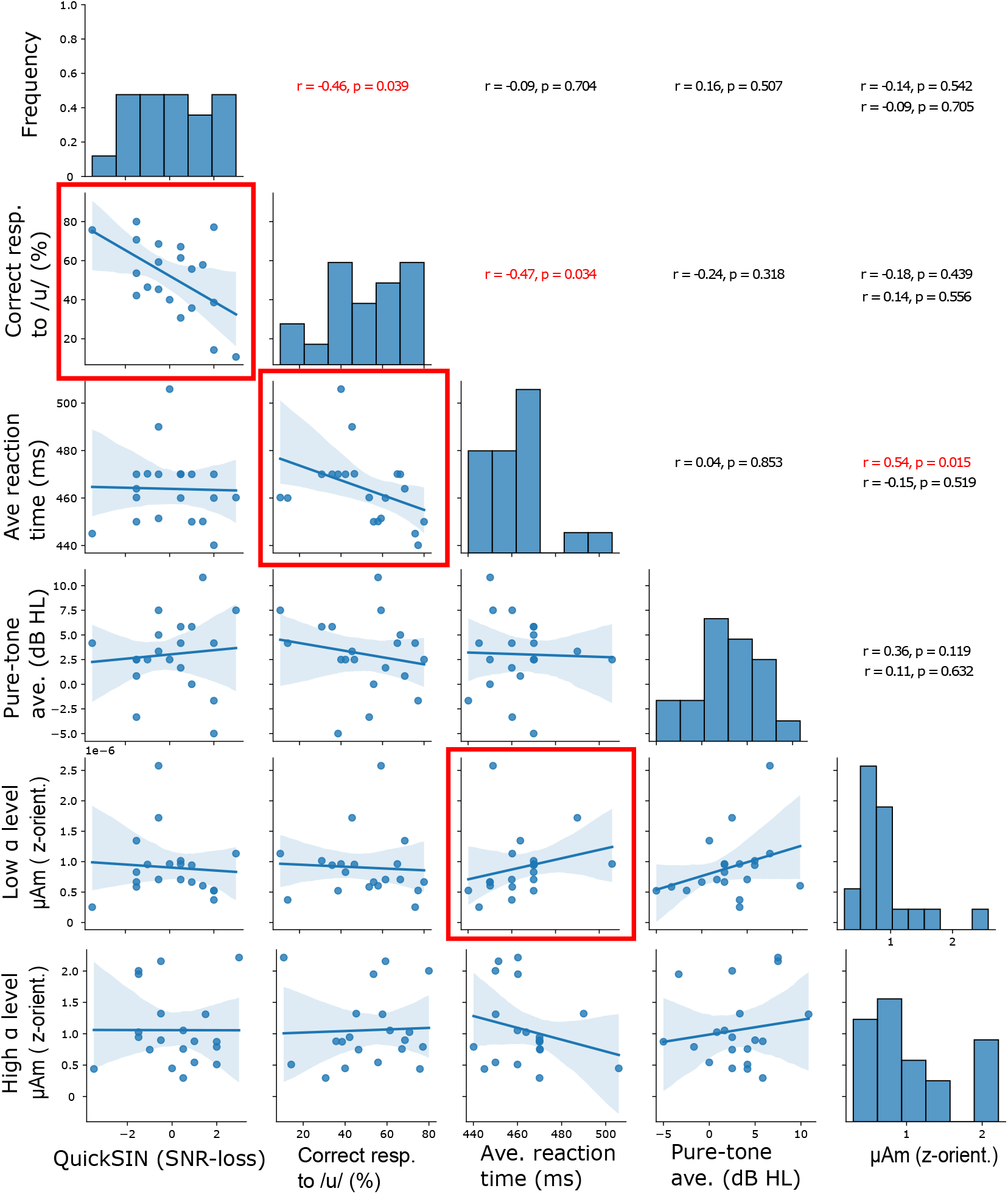
Summary of all correlations between pre-test perceptual performance vs. EEG behavioral task and EEG behavioral task vs. FFR F0 amplitudes (low or high *α* power). Indirect relationships between FFR F0 amplitudes at low/high *α* power and behavioral auditory performance are observed in the noise active condition. Correlations that are statistically significant (p < 0.05) are highlighted with red square boxes, and the corresponding r and p-values are shown in red text. The text in the last column show the r and p-values for low (upper) and high (lower) *α*, respectively. The distribution of the behavioral and brainstem response variables are shown along the diagonal. r = Spearman’s correlation, and shaded area indicates 95 % confidence intervals of the regression line.

## CRediT authorship contribution statement

**Jesyin Lai:** Conceptualization, Methodology, Software, Writing - Original draft preparation. **Caitlin N. Price:** Experimental design, Data collection. **Gavin M. Bidelman:** Conceptualization, Experimental design, Data curation, Writing - Original draft preparation.

